# Structure of yeast RAVE bound to a partial V_1_ complex

**DOI:** 10.1101/2024.07.18.604153

**Authors:** Hanlin Wang, Maureen Tarsio, Patricia M. Kane, John L. Rubinstein

## Abstract

Vacuolar-type ATPases (V-ATPases) are membrane-embedded proton pumps that acidify intracellular compartments in almost all eukaryotic cells. Homologous with ATP synthases, these multi-subunit enzymes consist of a soluble catalytic V_1_ subcomplex and a membrane-embedded proton-translocating V_O_ subcomplex. The V_1_ and V_O_ subcomplexes can undergo reversible dissociation to regulate proton pumping, with reassociation of V_1_ and V_O_ requiring the protein complex known as RAVE (regulator of the <u>A</u>TPase of <u>v</u>acuoles and <u>e</u>ndosomes). In the yeast *Saccharomyces cerevisiae*, RAVE consists of subunits Rav1p, Rav2p, and Skp1p. We used electron cryomicroscopy (cryo-EM) to determine a structure of yeast RAVE bound to V_1_. In the structure, RAVE is a L-shaped complex with Rav2p pointing toward the membrane and Skp1p distant from both the membrane and V_1_. Only Rav1p interacts with V_1_, binding to a region of subunit A not found in the corresponding ATP synthase subunit. When bound to RAVE, V_1_ is in a rotational state suitable for binding the free V_O_ complex, but it is partially disrupted in the structure, missing five of its 16 subunits. Other than these missing subunits and the conformation of the inhibitory subunit H, the V_1_ complex with RAVE appears poised for reassembly with V_O_.

## Introduction

Vacuolar-type adenosine triphosphatases (V-ATPases) are large membrane-embedded protein complexes that acidify intracellular compartments, including endosomes, lysosomes, the trans Golgi network, and secretory vesicles (Forgac, 2007). The role of these enzymes in organellar acidification makes them essential for many cellular processes, such as post-translational modification, trafficking of proteins, and degradation of material in lysosomes (Hinton et al., 2009). In some specialized cells, such as osteoclasts (Qin et al., 2012) and kidney intercalated cells (Brown et al., 2009), V-ATPases are targeted to the plasma membrane where they acidify the extracellular environment.

V-ATPases function through a rotary catalytic mechanism that couples ATP hydrolysis in their soluble catalytic V_1_ region to proton translocation through their membrane-embedded V_O_ region (reviewed in (Vasanthakumar and Rubinstein, 2020)). The V_1_ region includes a hexamer of alternating A and B subunits, with ATP hydrolysis occurring at three of the six A-B interfaces. Energy released by ATP hydrolysis induces conformational changes that cause rotation of a central rotor (subunits D, F, and d) and the attached membrane-embedded c ring (subunits c_8_, c′, c″, and Voa1p in the yeast *Saccharomyces cerevisiae*) (Mazhab-Jafari et al., 2016; Roh et al., 2018; Zhao et al., 2015). Rotation of the c ring causes proton translocation at the interface of the ring and the C-terminal membrane-embedded domain of subunit a. Three heterodimers of subunits E and G form peripheral stalk structures that interact with the A and B subunits of V_1_, the soluble subunits C and H, and the cytosolic N-terminal domain of subunit a from V_O_. These peripheral stalks prevent subunit a, and the associated transmembrane subunits e and f, from rotating along with the c ring. Electron cryomicroscopy (cryo-EM) of V-ATPases showed that when the purified enzyme is frozen and imaged at rest, it adopts three main conformations, known as rotational ‘State 1’, ‘State 2’, and ‘State 3’(Abbas et al., 2020; Zhao et al., 2015).

In *S. cerevisiae*, the V_1_ complex is formed in the cytosol, while the V_O_ complex assembles in the endoplasmic reticulum membrane with the assistance of dedicated assembly factors Vma12p, Vma21p, and Vma22p (Graham et al., 1998; Hill and Stevens, 1994; Wang et al., 2023; Wang and Rubinstein, 2023). Following assembly, the V_1_ and V_O_ complexes can undergo reversible dissociation in response to environmental cues (Kane, 1995; Sumner et al., 1995), presumably to regulate proton pumping and conserve ATP. In *S. cerevisiae*, dissociation occurs when glucose is depleted in the growth medium (Kane, 1995). The V_O_ complex and V_1_ complex missing subunit C (V_1_ΔC) can be immunoprecipitated after as little as 5 min of acute glucose deprivation (Kane, 1995), while V_1_ including subunit C (‘complete’ V_1_) can be isolated from yeast where glucose was depleted by culturing for ∼40 h (Vasanthakumar et al., 2022).

Following dissociation, the isolated V_O_ complex adopts an inhibited conformation in rotational State 3 (Mazhab-Jafari et al., 2016; Roh et al., 2018; Vasanthakumar et al., 2019) that is impermeable to protons (Qi and Forgac, 2008). The complete V_1_ complex is found in rotational State 2 while V_1_ΔC can adopt all three rotational states (Vasanthakumar et al., 2022). Cryo-EM of complete V_1_ and V_1_ΔC showed that subunit H forms an inhibitory interaction with one of the peripheral stalks, preventing ATP hydrolysis by the complexes (Parra et al., 2000; Vasanthakumar et al., 2022). However, X-ray crystallography of V_1_ΔC (Oot et al., 2016), cryo-EM of a preparation of V_1_ΔC that had been left at 4°C for several days (Vasanthakumar et al., 2022), and cryo-EM of intact lemon V-ATPase (Tan et al., 2022b) showed an alternative conformation of subunit H where it binds a subunit AB pair. Together, these observations suggest that subunit H has affinity for multiple surfaces on V_1_.

In yeast, both the initial assembly of V_1_ with V_O_ and their reassembly following dissociation require the RAVE complex (regulator of the <u>A</u>TPase of <u>v</u>acuoles and <u>e</u>ndosomes) (Jaskolka et al., 2021a; Seol et al., 2001; Smardon et al., 2002). RAVE is a soluble complex composed of subunits Rav1p, Rav2p, and Skp1p. In higher eukaryotes, the Rabconnectin-3 complex is functionally equivalent to RAVE and consists of at least Rabconnectin-3α (encoded by the genes *DMXL1* and *DMXL2*) and Rabconnectin-3β (encoded by *WDR7*) (Fagerberg et al., 2014; Kawabe et al., 2003). While Rabconnectin-3α has sequence homology with three regions of Rav1p, Rabconnectin-3β has no detectable homology with the subunits of RAVE (Jaskolka et al., 2021b). However, like Rav1p, both Rabconnectin-3α and β have been modelled to have N-terminal β propellers followed by α solenoid structures (Jumper et al., 2021). The individual contributions of Rabconnectin-3α and β to V-ATPase interactions and assembly are not known. Recent evidence suggests that Rogdi, which resembles yeast Rav2p, is also part of the Rabconnectin-3 complex (Winkley et al., 2021). In contrast, Skp1 has not been found in the Rabconnectin-3 (Cho et al., 2022; Jaskolka et al., 2021b).

RAVE associates with vacuolar membranes when glucose is present in the yeast growth medium and becomes predominantly cytosolic during glucose starvation (Smardon et al., 2015). The complex interacts with subunits E, G, and C of the V_1_ complex and the cytosolic N-terminal domain of the Vph1p-isoform of subunit a from the V_O_ complex. These interactions likely allow RAVE to facilitate V-ATPase reassembly by bringing V_1_ΔC, V_O_, and subunit C into close proximity (Jaskolka et al., 2021a; Jaskolka and Kane, 2020). However, *in vitro,* RAVE is not able to form a functional V-ATPase from V_1_ΔC, V_O_, and subunit C (Jaskolka et al., 2021a; Sharma et al., 2019). Reassembly can occur, and is accelerated by RAVE, when subunit H in V_1_ΔC is replaced with a non-inhibitory yeast-human chimeric subunit H.

We isolated a complex of yeast RAVE bound to V_1_ΔC and used cryo-EM to determine its structure. The structure shows that the three subunits of RAVE form an L-shaped complex that binds to a V_1_ΔC complex lacking a catalytic AB pair and peripheral stalk #3 (PS3). Beyond the loss of the AB pair and PS3, V_1_ΔC in the structure does not show substantial conformational changes compared to V_1_ΔC alone. The binding interface between RAVE and V_1_ΔC is formed by Rav1p and an area of the catalytic subunit A not found in the equivalent catalytic subunit of ATP synthase, and consequently known as the ‘non-homologous region’. V_1_ΔC in the structure is found in rotational State 3, which matches the rotational state of the dissociated V_O_ complex. Subunit H in the RAVE:V_1_ΔC complex remains in its inhibited conformation. The present structure suggests how RAVE interacts with isolated V_1_ following V-ATPase dissociation, and the events that need to occur before V_1_ and V_O_ can reassemble into an active V-ATPase.

## Results

### Overexpression of RAVE allows structure determination of a RAVE:V_1_ΔC complex

The structures of RAVE and RAVE interacting with V-ATPase have remained unknown, in part because of the low abundance of the endogenous RAVE complex (Breker et al., 2014; Jaskolka et al., 2021a). Overexpressing Rav1p is lethal to yeast, as it sequesters Skp1p from other essential protein complexes (Brace et al., 2006; Jaskolka et al., 2021a), which complicates the process of obtaining sufficient sample for structural analysis. To overcome these problems, we used a yeast strain that was engineered for galactose-induced overexpression of Rav1p and Rav2p-FLAG (Jaskolka et al., 2021a). Briefly, this strain was cultured in low glucose medium until the glucose was exhausted. Galactose was then added to induce Rav1p and Rav2p overexpression. In this condition, cell growth arrests after a single doubling, likely because of Skp1p depletion. Following this doubling, cells were harvested, lysed, and their cytosolic fraction isolated. RAVE was purified by anti-FLAG affinity chromatography making use of the FLAG tagged Rav2p. This method provides a complex of RAVE bound to V_1_ΔC (Jaskolka et al., 2021a). However, initial cryo-EM experiments showed disruption of the sample on grid freezing. Previous cryo-EM with V_1_ found that the complex is easily disrupted at the air-water interface of specimen grids, but this effect could be limited by addition of detergent to the sample (Vasanthakumar et al., 2022). Therefore, the sample was mixed with the detergent IGEPAL, frozen on cryo-EM specimen grids, and subjected to cryo-EM and image analysis (**Fig. S1** and **S2**). This analysis allowed calculation of a 3D map that shows RAVE bound to V1ΔC, although missing a catalytic AB pair and PS3 (**Fig. 1A**). Despite exploring numerous grid freezing conditions and methods, we were not able to obtain a map that included these missing subunits. Therefore, we went on to analyze the map and designate this complex RAVE:V_1_ΔC. The map had a nominal overall resolution of 2.7 Å, with local resolutions ranging from 2.4 to 7.8 Å (**Fig. S1C**, *blue curve*; **Fig. S2A**). The region of the map corresponding to RAVE was improved to a nominal overall 3.2 Å resolution with local refinement, with local resolutions ranging from 2.5 to 8.7 Å (**Fig. S1C**, *green curve*; **Fig. S2B**). Through fitting and refinement of an atomic model for V_1_ΔC (Vasanthakumar et al., 2022) and AlphaFold models for Rav1p, Rav2p, and Skp1p (Jumper et al., 2021), the map allowed for construction of an atomic model of the RAVE:V_1_ΔC complex (**Fig. 1B**, **Table S1**).

**Figure 1.**
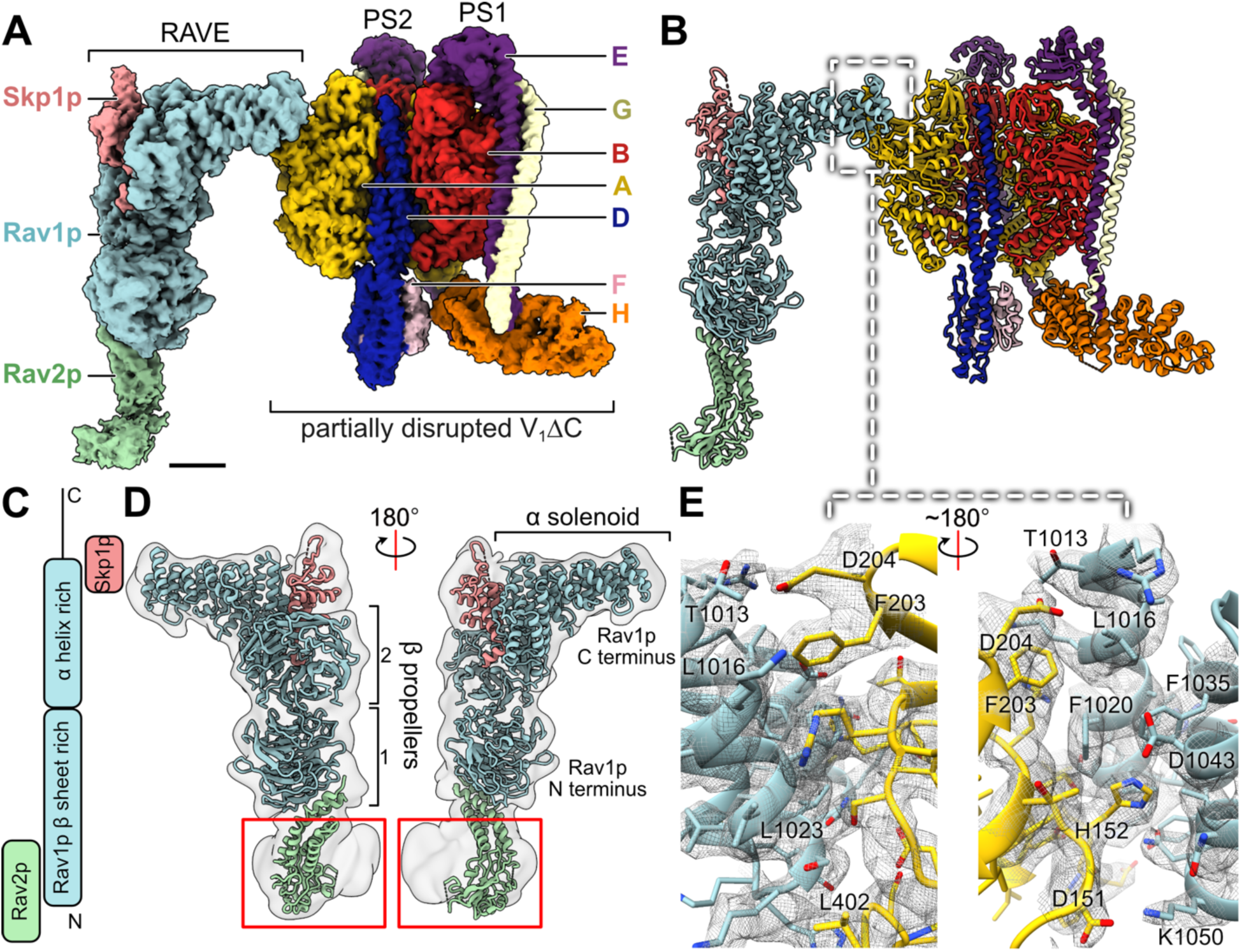
Structure of the RAVE:V_1_ΔC complex. **A**, Cryo-EM map of the RAVE:V**_1_**ΔC complex. Scale bar, 25 Å. **B,** Atomic model for RAVE:V_1_ΔC. **C,** Schematic representation of Rav1p domain organization and interactions. **D,** Fitting of RAVE from the RAVE:V_1_ΔC model in a low resolution map of RAVE alone. **E,** Amino acid side chains involved in the interaction between Rav1p from RAVE and subunit A from V_1_.

### Rav1p, Rav2p, and Skp1p assemble into an L-shaped complex

In the structure, Rav1p from RAVE serves as a scaffold for Rav2p and Skp1p. As determined biochemically, the N-terminal sequence of Rav1p interacts with Rav2p while its C-terminal sequence contacts Skp1p (Smardon et al., 2015) (**Fig. 1C and D**). Rav1p consists primarily of two N-terminal β propellers (residues 1-725) followed by an α solenoid (residues 762-1194) and a disordered C-terminal sequence (residues 1194 to 1357) that was not resolved in the structure (Jaskolka et al., 2021b; Kane and Smardon, 2003) (**Fig. 1C and D**). β propellers are composed of ‘blades’ of antiparallel β sheets and are often found to facilitate protein-protein interactions (Schapira et al., 2017; Xu and Min, 2011). The two β propellers in Rav1p are nearly perpendicular with each other and each has seven blades (**Fig. 1D**, *light blue*; **Fig. S3**). Within Rav1p, the α solenoid contacts the second β propeller, giving RAVE its L-shaped structure.

Rav2p consists primarily of four α helices and a β-sheet rich region near the bottom of the RAVE structure (**Fig. 1D**, *light green*; **Fig. S4A**). Residue 60 to 77, and 227 to 293 near the β-sheet rich region could not be built in the atomic model owing to low-resolution in this part of the map (**Fig. 1D**, *red rectangles*). The N and C termini from Rav2p interact with blades 3 and 4 of β propeller 1 in Rav1p (**Fig. S3**, *red rectangle*). The proposed human homologue of Rav2p, Rogdi (Cho et al., 2022; Winkley et al., 2021), is 7.7 kDa smaller than Rav2p but its crystal structure (Lee et al., 2017) aligns well with the structure of Rav2p determined here (**Fig. S4B**). Moreover, Rogdi was recently found to be enriched along with other Rabconnectin-3 subunits in a synaptic vesicle preparation from rat brain (Coupland et al., 2024). These findings provide further support that Rogdi is equivalent to Rav2p in mammalian cells and is a subunit of Rabconnectin-3.

### The structure suggests why Skp1p is largely dispensable for RAVE activity

Skp1p is a 22.3 kDa, 194 residue soluble protein, but only 127 residues were built in the RAVE:V_1_ΔC model (**Fig. 1D**, *light pink*; **Fig. S4C,** *left*). In the structure, Skp1p is mainly composed of five α helices and binds Rav1p between its α solenoid domain and its second β propeller domain (**Fig. 1A to D**, *light pink*). Skp1p is highly conserved in eukaryotes and is found in several different protein complexes (Connelly and Hieter, 1996; Orlicky et al., 2003; Seol et al., 2001). However, there is no evidence that mammalian Skp1 is part of the Rabconnectin-3 complex (Jaskolka et al., 2021b). Examination of the surface of Skp1p that binds Rav1p shows several Leu, Ile, Val, and Tyr residues that form a hydrophobic patch, which is also found in mammalian Skp1 (**Fig. S4C**, *orange surfaces*). This surface appears to be important for facilitating the multiple protein-protein interactions formed by both Skp1p and mammalian Skp1(Guan et al., 2021; Orlicky et al., 2003; Zheng et al., 2002) (**Fig. S4D**). Previously, the mutation of Asn139 in yeast Skp1p (corresponding to Asn108 in human Skp1) to Tyr was found to dissociate the protein from yeast RAVE but not the yeast SCF ubiquitin ligase (Brace et al., 2006). Asn139 is inside the binding site of Skp1p and Rav1p from RAVE, while slightly outside the binding site of Skp1p and Cdc4p from the SCF ubiquitin ligase (**Fig. S4E**), explaining why the mutation of this residue to a bulky Tyr can disrupt one interaction but not the other. In the RAVE:V_1_ΔC complex, Skp1p is far from V_1_, consistent with previous evidence that it is not required for V-ATPase assembly (Brace et al., 2006; Jaskolka et al., 2021b).

### The conformation of RAVE does not change on binding V_1_ΔC

We collected a second dataset of images with the detergent CHAPS added instead of IGEPAL, which gave rise to a lower-quality 3D map of RAVE alone (**Fig. S5** and **S6**). Although this map had a nominal overall resolution of 3.5 Å (**Fig. S5C**), the map suffered from anisotropic resolution owing to preferred orientation of the particle images (**Fig. S6B** and **C**) and was filtered to 12 Å, which was the resolution of the worst direction in the anisotropic map (**Fig. 1D**, *semi-transparent gray surface***; Table S1**). Despite the low resolution of the map, fitting the atomic model of RAVE from the RAVE:V_1_ΔC structure into the map of RAVE alone (**Fig. 1D**) showed no obvious conformational difference between the two structures. This finding indicates that RAVE does not undergo major conformational changes upon binding V_1_.

### Only Rav1p from RAVE interacts with V_1_ΔC and binding does not change V_1_ΔC conformation

The interaction between Rav1p and V_1_ΔC is mediated solely by three α helices (residues 1012-1024, 1028-1037, and 1041-1050) from Rav1p’s α solenoid domain, which bind subunit A in V_1_ (**Fig. 1E**). These α helices are in the region of Rav1p (residues 840-1150) with the highest sequence similarity to Rabconnectin-3α (Jaskolka et al., 2021b). The interaction site appears to be stabilized by van der Waals interactions involving Leu1016, Phe1020, Leu1023, Phe1035 from Rav1p, and Phe203 and Leu402 from subunit A, and possible hydrogen bonds between Asp1043, Thr1013, Lys1050 from Rav1p and His152, Asp204, Asp151 from subunit A, respectively (**Fig. 1E**).

To examine if binding of RAVE induces conformational changes in V_1_, we fit the atomic model of V_1_ΔC from the RAVE:V_1_ΔC structure into previously published maps of V_1_ΔC (Vasanthakumar et al., 2022) (**Fig. 2A**, **Fig. S7**). This fitting shows that the partially disrupted V_1_ΔC from the RAVE:V_1_ΔC structure is in rotational State 3, and that there is no substantial conformational change when RAVE binds V_1_ΔC. Interestingly, the RAVE:V_1_ΔC map shows clear density for an additional ADP molecule in one catalytic AB pair that was not built in the crystal structure of V_1_ΔC and was not resolved in the cryo-EM map of V_1_ΔC (Oot et al., 2016; Vasanthakumar et al., 2022) (**Fig. S8A**). The role of this ADP molecule is unclear. The consistent V_1_ΔC conformation, whether alone or when bound to RAVE, allowed us to consider what other interactions would occur between Rav1p and V_1_ΔC if V_1_ΔC were intact. This comparison suggests that the missing subunit B and subunits E and G from PS3 would contact Rav1p if they were present (**Fig. 2B**, *semi-transparent surfaces*). These interactions would be consistent with the finding that subunits E and G are important for RAVE binding to V_1_ (Smardon et al., 2002).

**Figure 2.**
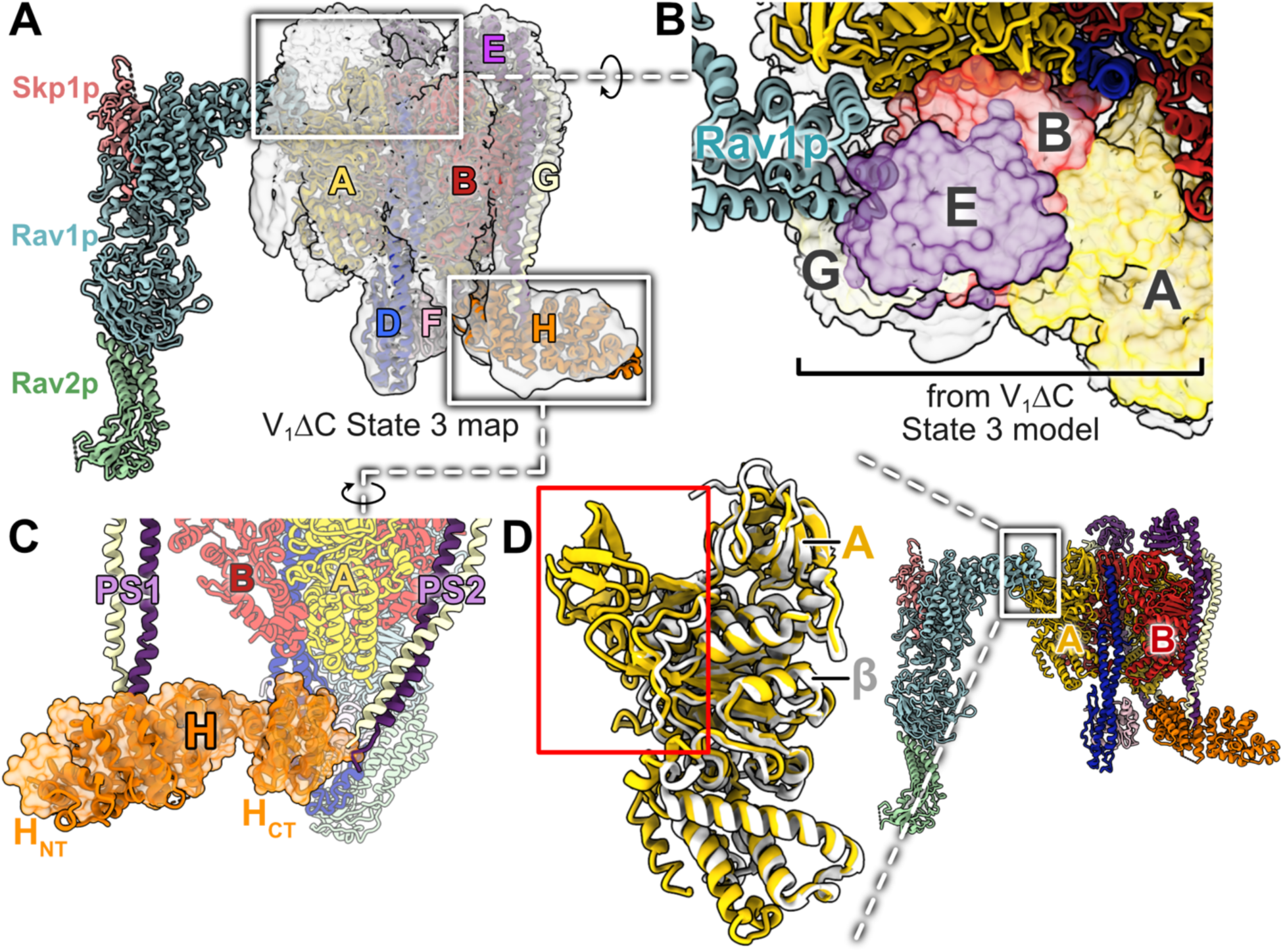
Characterization of the partially disrupted V_1_ΔC in the RAVE:V_1_ΔC complex. **A**, Fitting of the RAVE:V_1_ΔC complex atomic model in a density map of intact V_1_ΔC State 3 (EMD-25999). **B,** Addition of the missing V_1_ subunits from V_1_ΔC in State 3 (semi-transparent surfaces; from PDB 7TMQ) to the RAVE:V_1_ΔC model. **C,** The RAVE:V_1_ΔC structure shows that subunit H forms the same inhibitory interaction with PS2 seen in structures of complete V_1_ and V_1_ΔC in all three rotational states. The semi-transparent model of subunit H (orange) is from V_1_ΔC State 3 (PDB 7TMQ). **D,** Structural alignment of subunit A of yeast V-ATPase (yellow) with subunit β of yeast ATP synthase (white; PDB 7TKO). The non-homologous region of V-ATPase subunit A is highlighted with a red rectangle.

In the RAVE:V_1_ΔC structure, subunit H adopts the ATP hydrolysis-inhibiting conformation seen previously in both complete V_1_ and V_1_ΔC (**Fig. 2C**) (Vasanthakumar et al., 2022). In this conformation, the N-terminal domain of subunit H (H_NT_) remains bound to peripheral stalk #1 (PS1), as it is in the intact V-ATPase structure, while the C-terminal domain (H_CT_) binds a site on peripheral stalk #2 (PS2). This linking of PS1 and PS2 pulls PS2 across a catalytic AB pair, inhibiting the enzyme. The distance between RAVE and the inhibitory subunit H in the RAVE:V_1_ΔC structure suggests that RAVE cannot alter the conformation of H, explaining why RAVE alone cannot restore ATP hydrolysis when mixed with wildtype V_1_, V_O_, and subunit C (Jaskolka et al., 2021a).

### RAVE binds the non-homologous region of subunit A

V-ATPases are structurally and evolutionarily related to the F-type ATP synthases found in bacteria, mitochondria, and chloroplasts (reviewed in (Courbon and Rubinstein, 2022)). The catalytic subunit A from V_1_ is homologous with subunit β from the ATP synthase F_1_ region (Bowman et al., 1988; Guo and Rubinstein, 2022; Zimniak et al., 1988) (**Fig. 2D**, *left*). However, there is a conserved ∼90 residue sequence in subunit A that is not present in the β subunit, which is known as the ‘non-homologous region’ (Shao et al., 2003) (**Fig. 2D**, *red rectangle*). Notably, the RAVE:V_1_ΔC structure shows that Rav1p binds to the non-homologous region of subunit A.

### The conformation of V_1_ΔC selects for RAVE binding to a single site in rotational State 3

There are three catalytic interfaces between subunits A and B in V_1_, known as the ‘open’, ‘occluded’, and ‘ADP-bound’ interfaces (Abbas et al., 2020; Zhao et al., 2015) (**Fig. 3A**, *blue curves*). There are also three non-catalytic A-B interfaces. If the V_1_ΔC complex in the RAVE:V_1_ΔC structure were intact, RAVE would be found at a non-catalytic interface (**Fig. 3A**). Interestingly, although three nearly equivalent binding sites in the V_1_ region exist at each of the non-catalytic interfaces, the structure shows RAVE only bound to the site near the missing PS3 (**Fig. 3A i**). Adding the missing PS3 from the published structure of V_1_ΔC in rotational State 3 to the RAVE:V_1_ΔC model causes only minimal clashes (**Fig. 3A i**, *red circle*). In contrast, RAVE appears to be prevented from binding the other two non-catalytic interfaces by differences in the conformations of these sites. Compared to PS3, PS1 and PS2 are bent toward each other because of their interactions with subunit H (Vasanthakumar et al., 2022) (**Fig. 2C**). This bending of the peripheral stalks reduces the space available for RAVE to bind the non-catalytic interfaces near PS1 and PS2, which appears to be sufficient to prevent binding (**Fig. 3A ii** and **iii**). As described above, RAVE binds V_1_ΔC only in rotational State 3. In rotational State 3, the conformation of PS3 provides a sufficiently large binding site for RAVE (**Fig. 3B i**). However, aligning PS3 from V_1_ΔC complexes in all three rotational states shows that PS3 is closer to subunit A in State 1 and 2, than it is in State 3, which prevents RAVE from binding these states (**Fig. 3B ii** and **iii**).

**Figure 3.**
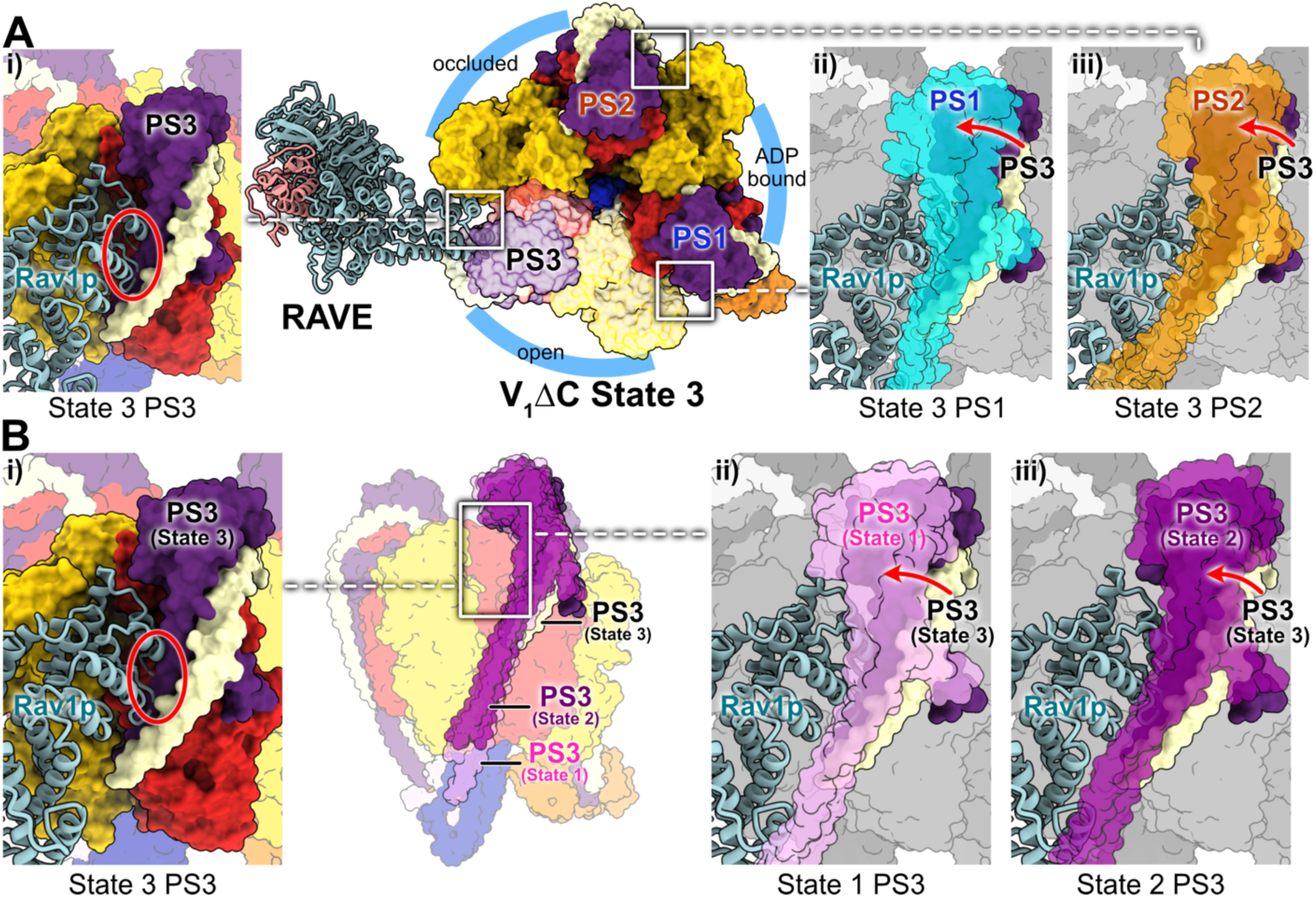
Binding site of RAVE on subunit A. **A**, Mapping of Rav1p onto its observed binding site near the location of PS3 (**i**, purple and beige surfaces) versus the non-catalytic AB interfaces near PS1 (**ii**, cyan surface) and PS2 (**iii**, orange surface). **B,** Mapping of Rav1p binding onto its observed site near the location of PS3 in rotational State 3 (**i**, purple and beige surfaces) versus rotational State 1 (**ii**, pink surface; from PDB 7TMO) and rotational State 2 (**iii**, violet surface; from PDB 7TMP).

### The observed conformation of RAVE holds V_1_ΔC above the membrane

Cryo-EM of the isolated V_O_ complex has consistently found the complex in rotational State 3 (Keon et al., 2022; Mazhab-Jafari et al., 2016; Roh et al., 2018; Vasanthakumar et al., 2019). The observation that RAVE binds V_1_ΔC in rotational State 3 suggests that, other than the inhibitory conformation of subunit H, the absence of subunit C, and possibly binding of inhibitory ADP, RAVE:V_1_ΔC and V_O_ are compatible for reassembly. However, when Rav2p in RAVE is brought into contact with the expected position of the membrane, subunit D from V_1_ΔC is at least ∼15 Å above subunit d from V_O_ (**Fig. 4A and B**, *gray dashed line*). This difference in height indicates that a conformational change must occur in RAVE, or the membrane must bend, for V_1_ΔC to come close enough to V_O_ for the V-ATPase to assemble.

**Figure 4.**
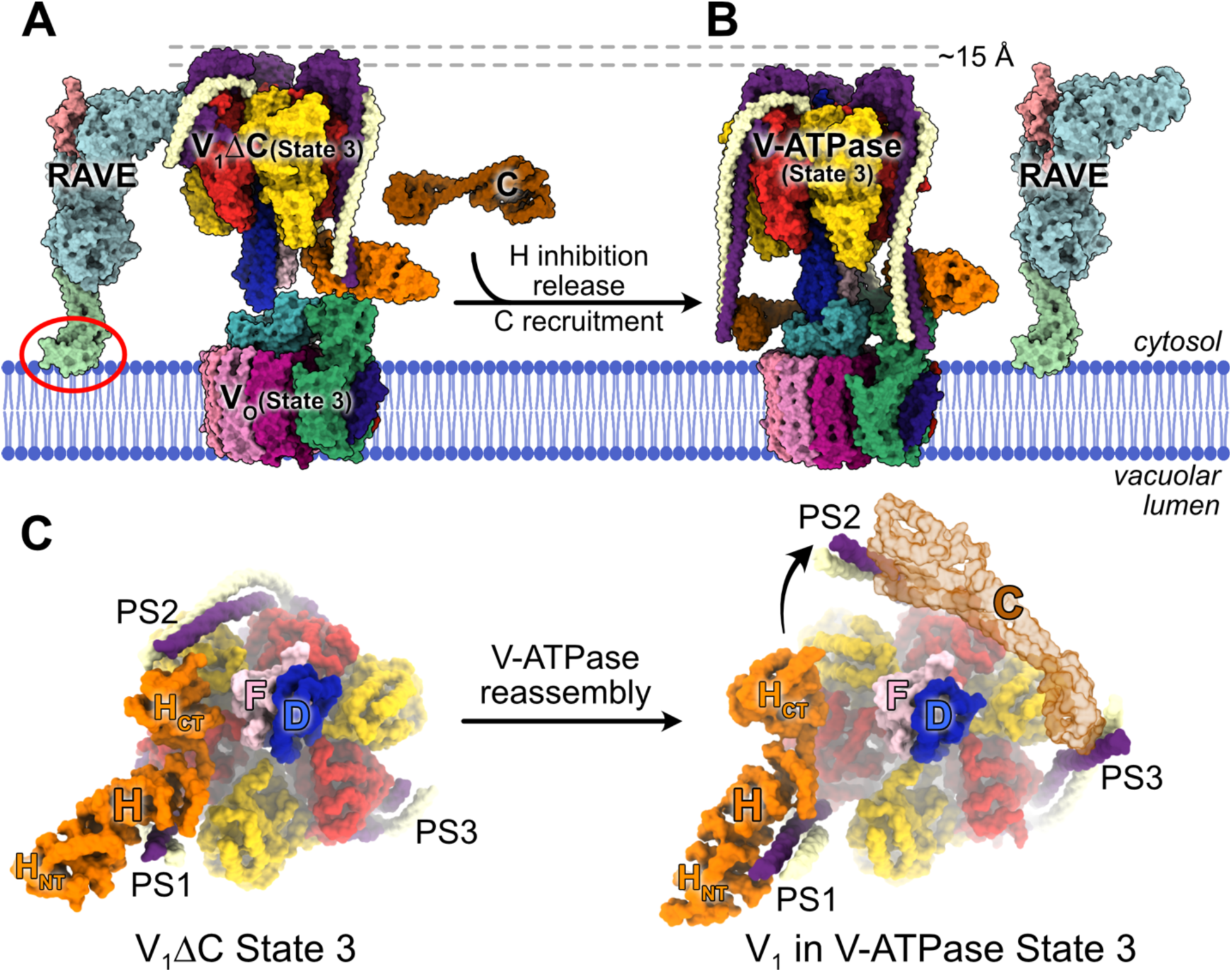
Model for V-ATPase reassembly by RAVE. **A**, RAVE brings V_1_ΔC close to the isolated V_O_, but a clash between Rav2p and the membrane (red circle) leaves V_1_ above its binding site in V_O_. **B,** After return of subunit H to its non-inhibitory conformation and recruitment of subunit C, V-ATPase reassembly releases the RAVE complex. **C,** During V-ATPase reassembly, subunit H is released from its inhibitory conformation, allowing PS2 and PS3 to bind subunit C (transparent brown subunit) as seen in the intact V-ATPase State 3 (PDB 7TMT).

## Discussion

The RAVE:V_1_ΔC structure reported here shows only limited contacts between RAVE and V_1_. Fitting of a complete model for V_1_ΔC into the RAVE:V_1_ΔC map indicates that in the intact structure, Rav1p would bind subunits B, E, and G, as well as A (**Fig. 2B**). The overall interaction area is still surprisingly small given RAVE’s role in reattaching V_1_ to V_O_. However, the region of Rav1p that interacts with subunit A is highly conserved in Rabconnectin-3α, supporting its functional importance. The bulk of Rav1p consists of β propeller structures, which often form protein-protein interactions (Schapira et al., 2017; Xu and Min, 2011). The first β propeller domain of Rav1p, which binds Rav2p, is another region that is conserved in Rabconnectin-3α (Jaskolka et al., 2021b; Smardon et al., 2015). Although Skp1p appears dispensable for RAVE function (Brace et al., 2006), loss of Rav1p or Rav2p results in defects in cell growth and V-ATPase assembly (Seol et al., 2001; Smardon et al., 2002). However, like Skp1p, Rav2p does not interact with V_1_ in the structure, and consequently the molecular function of Rav2p in reassembly remains unclear.

Binding of Rav1p to the non-homologous region of subunit A suggests that this region is important in reversible dissociation of V-ATPase, which is a phenomenon that does not occur with ATP synthases (Kane, 1995). Interestingly, the isolated non-homologous region of subunit A was found to interact with V_O_ in a glucose-dependent manner (Shao and Forgac, 2004). The interaction between RAVE and the non-homologous region could allow RAVE to link this region to V_O_. The non-homologous region of subunit A also interacts with a *Legionella pneumophila* effector protein SidK, which inhibits V-ATPase by ∼30% (**Fig. S8B**, *cross-sections*) (Maxson et al., 2022; Zhao et al., 2017). SidK binds V-ATPase with high affinity (Sharma and Wilkens, 2017; Xu et al., 2010; Zhao et al., 2017), a property that has allowed the protein to be used to isolate both V-ATPase and intact synaptic vesicles (Abbas et al., 2020; Coupland et al., 2024; Tan et al., 2022b, 2022a), and to localize V-ATPase in fluorescence microscopy images (Maxson et al., 2022). The shared binding site for RAVE and SidK could result in SidK binding interfering with RAVE-mediated V-ATPase assembly. This observation may help explain why SidK overexpression in mammalian cells produces an unusual phenotype, with lysosome pH increased by only ∼0.25 but the lysosomes localizing to the cell periphery, which is characteristic of potent V-ATPase inhibition (Maxson et al., 2022).

The RAVE:V_1_ΔC structure suggests how RAVE associates with V_1_ΔC when glucose is depleted in the yeast growth medium. When glucose is restored, several steps need to occur before RAVE can reassemble functional V-ATPase from V_1_ΔC, subunit C, and V_O_. First, subunit H is found in the inhibitory conformation seen previously, with H_CT_ pulling PS2 across a catalytic AB interface (**Fig. 2C**; **Fig. 4C**, *left*) (Vasanthakumar et al., 2022). Therefore, for V-ATPase activity to resume, the connection between subunit H and PS2 must be broken. Release of PS2 from H_CT_ would allow PS2 to move further from the central stalk subunits D and F (**Fig. 4C**, *black arrow*), enabling subunit C to bind both PS2 and PS3, as observed in the intact V-ATPase (**Fig. 4C**, *right*). This required conformational change suggests that during reassembly, subunit H and PS2 must separate before subunit C can be recruited to V_1_ΔC.

The structure described here does not account for several reported interactions between RAVE and other V-ATPase subunits. It does not explain the observation that RAVE alone and RAVE:V_1_ΔC can bind to subunit C, the latter with higher affinity (Jaskolka et al., 2021a; Smardon and Kane, 2007). Further, it is not clear from the structure how Rav1p in RAVE would interact with the N-terminal domain of the subunit a isoform Vph1p (Smardon et al., 2015, 2014). These interactions suggests that other conformations of RAVE may exist. Consistent with this hypothesis, RAVE binds the same side of V_1_ΔC that subunit C would bind (**Fig. 4A and B**), which could facilitate interaction of RAVE and subunit C. For V-ATPase assembly, RAVE also needs to bring V_1_ closer to V_O_, with the lipid bilayer presenting an apparent barrier to that process (**Fig. 4A**, *red circle*). Notably, Rav2p has been shown to bind to subunit C (Smardon et al., 2015). A Rav2p-subunit C interaction could be part of a conformational change that both assists in placing subunit C in V_1_ and moves Rav2p further from the membrane, allowing access to V_O_. Therefore, the structure described here likely presents just one snapshot in the process of RAVE-mediated V-ATPase reassembly.

## Author contributions

HW purified the protein sample and performed cryo-EM and associated analysis. MT grew the yeast cultures. JLR and PMK conceived and supervised the research. HW and JLR wrote the manuscript and prepared figures, with input from the other authors.

## Acknowledgements

We thank Samir Benlekbir and Zhijie Li for assistance with cryo-EM data collection. HW was supported by an Ontario Graduate Scholarship and a Mary H. Beatty Fellowship. JLR was supported by the Canada Research Chairs program. This research was supported by Canadian Institutes of Health Research grant PJT166152 (JLR) and National Institutes of Health grant 1R35GM145256 (PMK). Cryo-EM data was collected at the Toronto High-Resolution High-Throughput cryo-EM facility, supported by the Canada Foundation for Innovation and Ontario Research Fund.

## Data availability

Electron microscopy maps are available from the electron microscopy databank with accession codes EMD-45788 and EMD-45789, and the atomic model is available from the protein databank with accession code 9COP.

## Methods

### Yeast Strain Construction and Yeast Growth

The haploid *S. cerevisiae* strains SF838-5Aα *P_GAL1_-RAV1* and SF838-5Aa *P_GAL1_-RAV2*-FLAG were crossed to create a diploid with both *RAV1* and *RAV2*-FLAG under control of the *GAL1* promoter as described (Jaskolka et al., 2021a). Briefly, to grow yeast for protein purification, a 1 L culture of the diploid cells was grown at 30°C with shaking to a density of 1 OD_600_/ml in YEPD (1% [w/v] yeast extract, 2% [w/v] peptone, 2% [w/v] dextrose medium) adjusted to pH 5 by addition of 40 ml of 1 M HCl. Cells were harvested by centrifugation at 3475 ×g (3500 rpm, Beckman JS4.2 rotor) and washed in YEP medium with no dextrose. The washed cells were inoculated into YEP containing 0.2% (w/v) glucose at a density of 0.05 OD_600_/ml and grown to a density of 1 OD_600_/ml over 16-18 h. Galactose was added to the cultures to give a final concentration of 2% (w/v) and growth was continued at 30°C for another 14 h. After this time, the cells had doubled once but growth had arrested. The cells were then collected by centrifugation as described above and washed with water, followed by a second wash in phosphate buffered saline (PBS; 137.9 mM NaCl, 2.67 mM KCl, 1.47 mM KH_2_PO_4_, 8 mM Na_2_HPO_4_, 2 mM EDTA, pH 7.4), and finally suspended in a volume of PBS equal to the volume of the cell pellet. Resuspended cells corresponding to 4000 OD_600_ units (2 L of final culture) were frozen in 50 ml centrifuge tubes at –80 °C.

### Protein Purification

Frozen cell pellets were thawed at room temperature, and all subsequent purification steps were performed at 4°C. Cells were resuspended in PBS lysis buffer (137.9 mM NaCl, 2.67 mM KCl, 1.47 mM KH_2_PO_4_, 8 mM Na_2_HPO_4_, 2 mM EDTA, 5 mM 6-aminocaproic acid, 5 mM benzamidine hydrochloride, 5 mM ethylenediaminetetraacetic acid [EDTA], 10 mg/L phenylmethylsulfonyl fluoride [PMSF], pH 7.4), and were lysed with 10 passages through an Avestin Emulsiflex C3 Homogenizer at 20,000 psi, with 5 min rest on ice between each passage. Cell debris was removed by centrifugation at 41,140 ×g for 45 min (17,000 rpm, Beckman Avanti, JA-20 rotor) and membranes were removed by ultracentrifugation at 396,000 ×g for 1.5 h (55,000 rpm, Beckman L-90K, Ti70 rotor). The supernatant was filtered with a 0.45 μm syringe filter and applied to 1.5 mL anti-FLAG M2 affinity gel (Millipore Sigma) in a disposable plastic column, pre-equilibrated in PBS (137.9 mM NaCl, 2.67 mM KCl, 1.47 mM KH_2_PO_4_, 8 mM Na_2_HPO_4_, 2 mM EDTA, pH 7.4). Material that flowed through the column was collected and applied to the column again. The column was washed with ten column volumes of PBS and protein was eluted with three column volumes of PBS containing 150 μg/mL of 3×FLAG peptide and one column volume of PBS without peptide. Protein was concentrated to ∼200 μL with a 100 kDa molecular weight cutoff (MWCO) Amicon Ultracentrifugal filter (Millipore Sigma) at 1000 ×g. Samples were further concentrated to ∼1-2 mg/mL with a 100 kDa MWCO Vivaspin 500 centrifugal concentrator (Sartorius) at 1000 ×g.

### Cryo-EM Specimen Preparation and Data Collection

IGEPAL CA-630 (Sigma) or 3-[(3-cholamidopropyl) dimethylammonio]-2-hydroxy-1-propanesulfonate (CHAPS; Anatrace) were added to final concentrations of 0.025% (v/v) and 0.1% (v/v), respectively. Samples were applied to homemade nanofabricated holey gold grids with regular arrays of ∼2 μm holes (Marr et al., 2014). Grids were glow-discharged for 2 min, and 1.5 μL of sample was applied to each grid, blotted for 2 s at 4 °C and 80% relative humidity, and frozen in liquid ethane with a Leica EM GP2 freezing device. Sample screening was performed with a FEI Tecnai F20 electron microscope operating at 200 kV and equipped with a Gatan K2 Summit direct detector device camera. High-resolution data collection was performed with a Titan Krios G3i electron microscope (Thermo Fisher Scientific) operating at 300 kV and equipped with a Falcon 4i camera. Data collection was automated with the *EPU* software package (Thermo Fisher Scientific). Electron event representation movies were collected (Guo et al., 2020) with a calibrated pixel size of 1.03 Å/pixel and a total exposure of ∼42 e^-^/Å^2^ for IGEPAL dataset and ∼40 e^-^/Å^2^ for CHAPS dataset, and fractionated into 30 exposure fractions.

### Image Analysis

Movie alignment with patch-based motion correction and estimation of contrast transfer function (CTF) parameters were performed with *cryoSPARC Live*. All other image analysis steps were performed with *cryoSPARC v3* (Punjani et al., 2017). After removing movies with poor CTF fit, ice thickness, or motion, 4,718 movies from the IGEPAL datasets and 5,341 movies from the CHAPS dataset were selected for further processing. Templates for particle selection were generated from 2D classification of manually selected particle images. Individual particle motion correction was performed (Rubinstein and Brubaker, 2015) and datasets were cleaned with multiple rounds of 2D classification and *ab initio* 3D classification. This process provided 66,347 and 214,437 particle images for Topaz training on IGEPAL and CHAPS datasets, respectively (Bepler et al., 2019). After multiple rounds of 2D classification and *ab initio* 3D classification for cleaning, 71,343 and 92,040 particles images were used for three-dimensional structure determination from the IGEPAL and CHAPS datasets, respectively.

For both the IGEPAL and CHAPS datasets, *ab initio* 3D classification and heterogeneous refinement were used to generate initial 3D maps of RAVE:V_1_ΔC and RAVE. For the IGEPAL dataset, the RAVE:V_1_ΔC map was refined with nonuniform refinement (Punjani et al., 2020), followed by two rounds of local and global CTF refinement, nonuniform refinement, and one round of reference-based motion correction. This analysis led to a map with an overall nominal resolution of 2.7 Å. A mask on V_1_ΔC was generated for RAVE:V_1_ΔC particle subtraction, and local refinement of the RAVE region of the map yielded a 3.2 Å resolution map of RAVE. For the CHAPS dataset, the RAVE map was refined with nonuniform refinement, followed by two rounds of local and global CTF refinement, and another round of nonuniform refinement. The RAVE map, which suffered from anisotropic resolution, was low-pass filtered to 12 Å resolution, which was the worst resolution direction in the map.

### Atomic model building

The previously published V_1_ model 7TMQ (Vasanthakumar et al., 2019), and AlphaFold models of Rav1p, Rav2p, and Skp1p (Jumper et al., 2021) were fit into the map of RAVE:V_1_ΔC with UCSF Chimera (Pettersen et al., 2004). These models were adjusted manually in *Coot* (Emsley and Cowtan, 2004), followed by refinement with ISOLDE (Croll, 2018), and real space refinement with PHENIX (Adams et al., 2010). Figures were rendered with Chimera (Pettersen et al., 2004) and UCSF ChimeraX (Goddard et al., 2018).

## Figures

**Figure S1.**
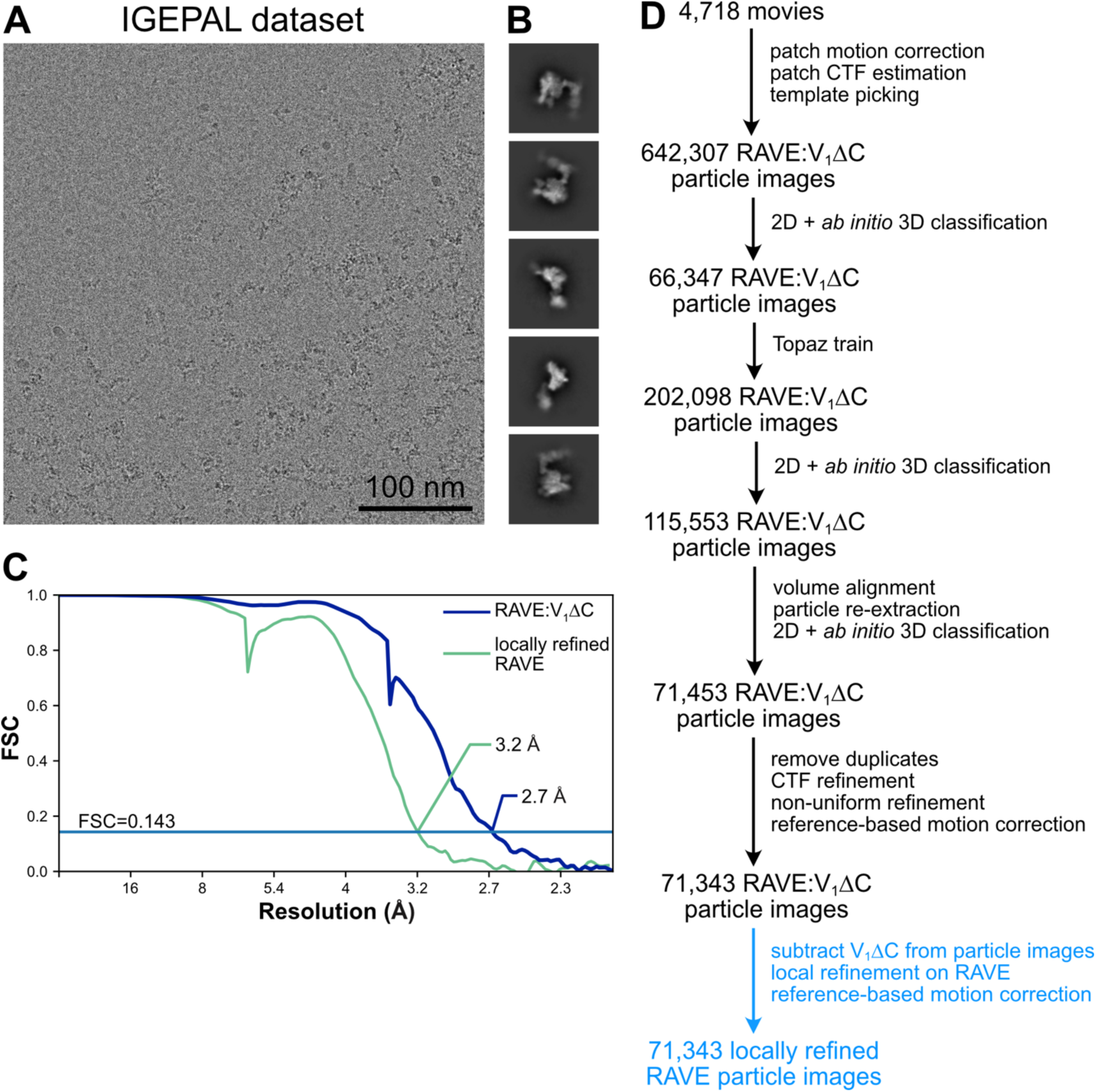
Cryo-EM of the RAVE:V_1_ΔC complex. Example micrograph (**A**) and 2D class average images (**B**). **C,** Fourier shell correlation curves, corrected for masking, after gold-standard refinement for RAVE:V_1_ΔC and locally refined RAVE. **D,** Workflow to obtain map of the RAVE:V_1_ΔC complex.

**Figure S2.**
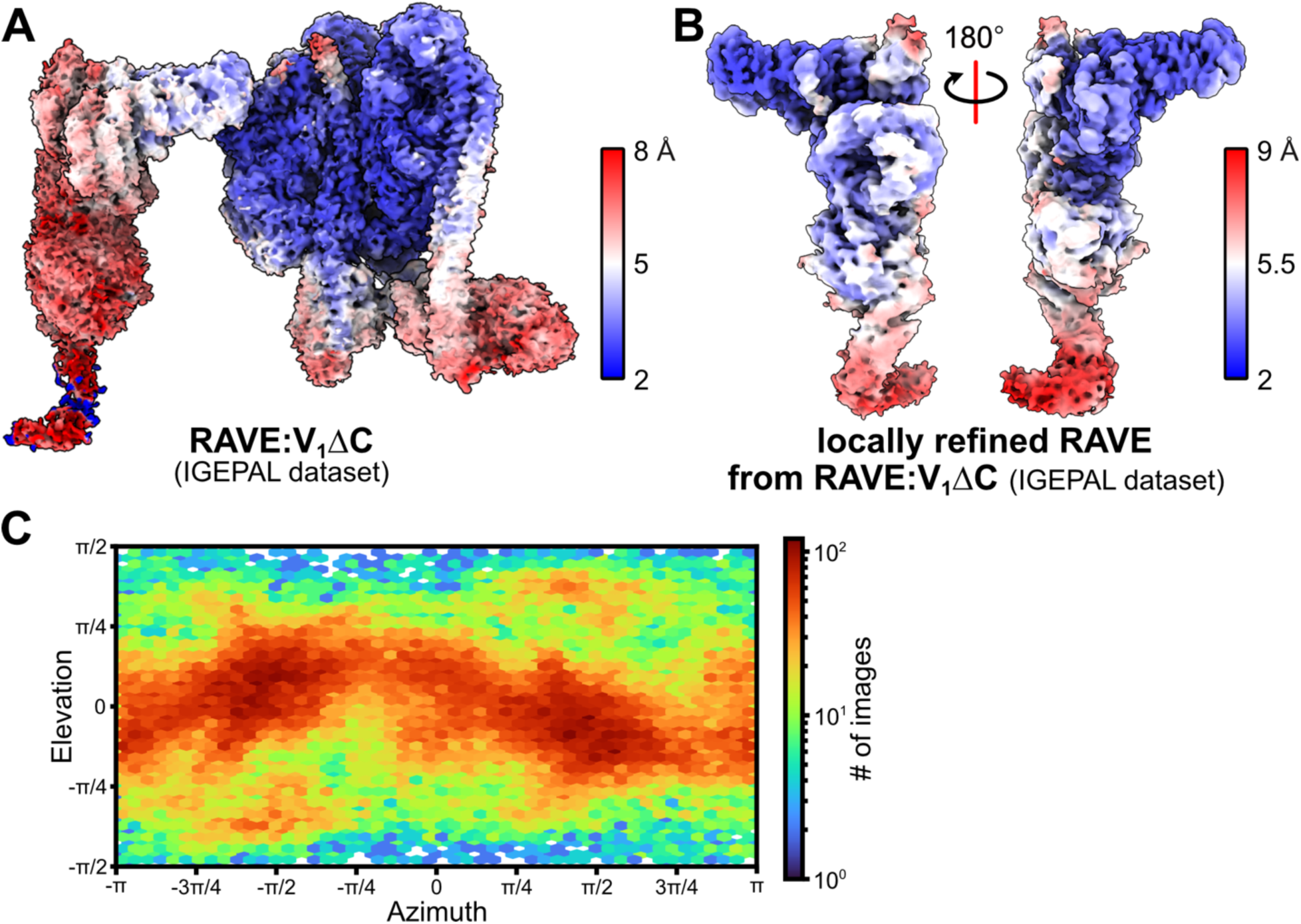
Local resolution and particle orientation distribution for the RAVE:V_1_ΔC complex. **A**, Local resolution map for RAVE:V_1_ΔC. **B,** Local resolution map for the locally refined RAVE region of the map. **C,** Orientation distribution plot for RAVE:V_1_ΔC.

**Figure S3.**
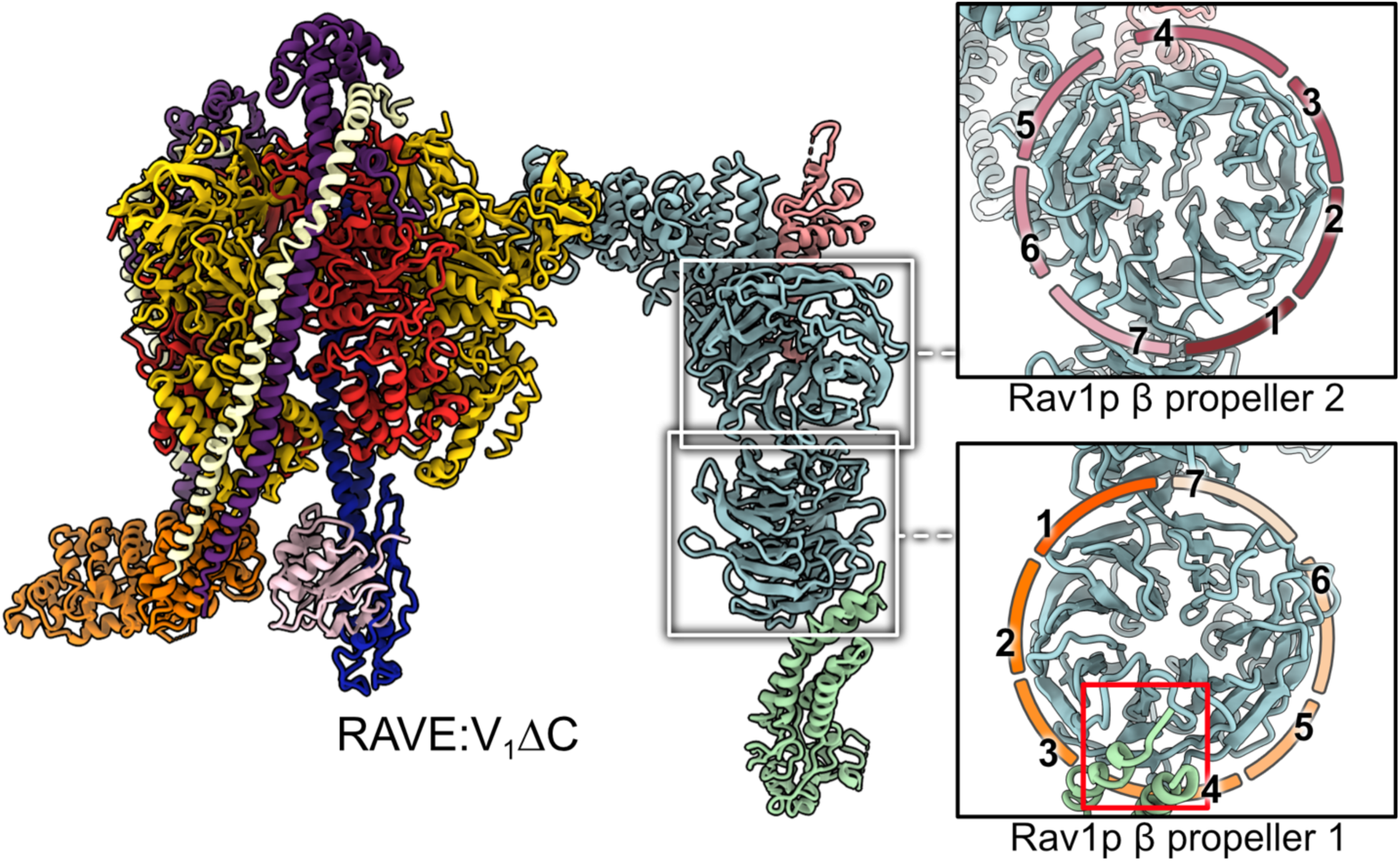
The β propeller structures of Rav1p in the RAVE:V_1_ΔC complex. The N-terminal domain of Rav1p is composed of two β propellers, each with seven ‘blades’. The interaction between blade 3 and 4 from Rav1p β propeller 1 and Rav2p is highlighted with a red rectangle.

**Figure S4.**
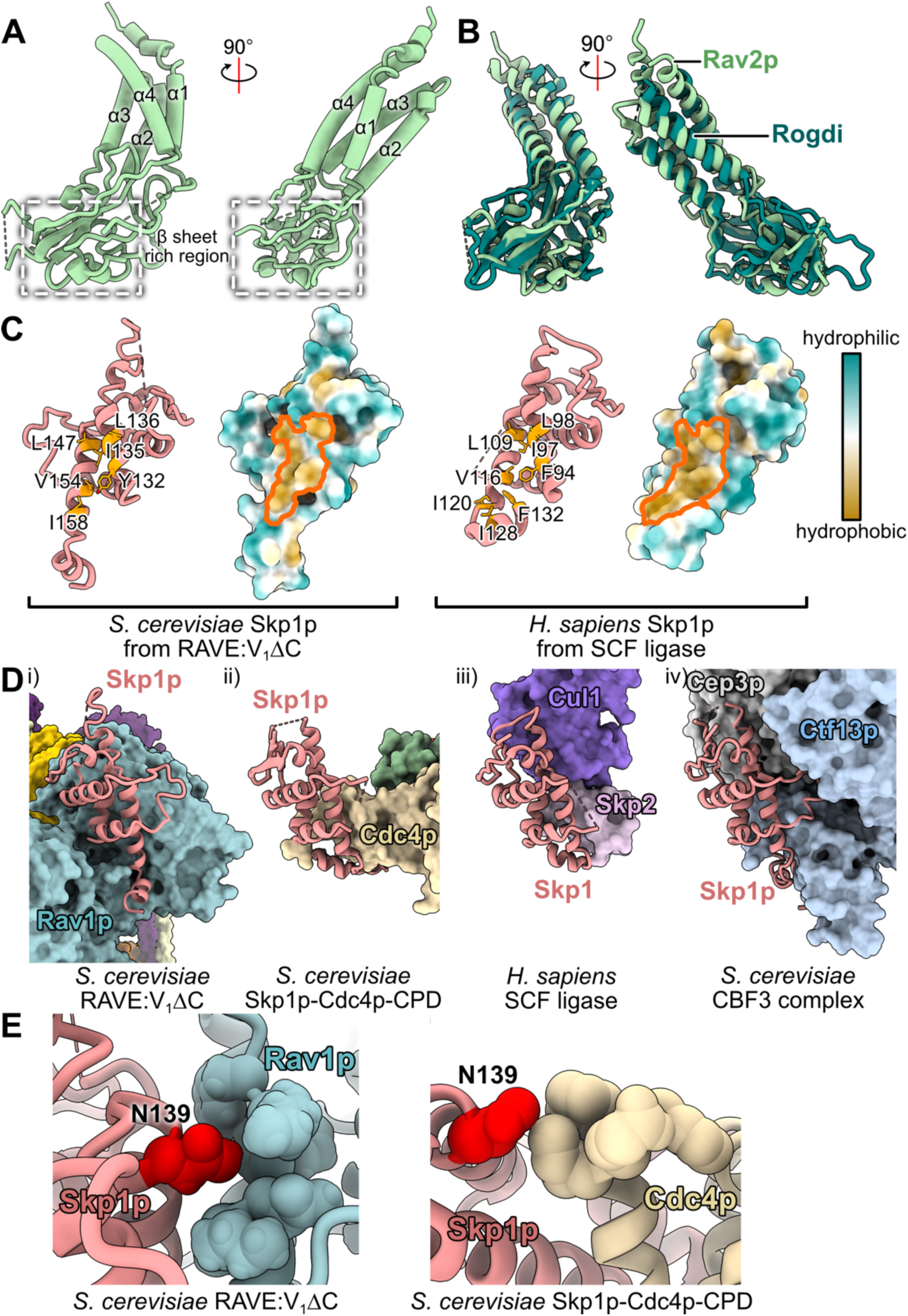
Structural features in Rav2p and Skp1p. **A**, Domain organization of Rav2p from the RAVE:V_1_ΔC complex. **B,** Structural alignment of Rav2p with the crystal structure of human Rogdi (PDB 5XQH). **C,** Surface hydrophobicity of yeast Skp1p and human Skp1 (from PDB 1LDK). Residues in the hydrophobic patches are highlighted in orange. **D,** Comparison of yeast Skp1p and human Skp1 interactions with their binding partners in RAVE:V_1_ΔC (**i**), yeast Skp1p-Cdc4p-CPD (**ii**; PDB 1NEX), human SCF uniquitin ligase (**iii**; PDB 1LDK), and the yeast CBF3 complex (**iv**; PDB 7K79). **E,** Asn139 in Skp1p is within the binding site of Skp1p and Rav1p from RAVE (*left*), but not Skp1p and Cdc4p from the SCF ubiquitin ligase (*right*).

**Figure S5.**
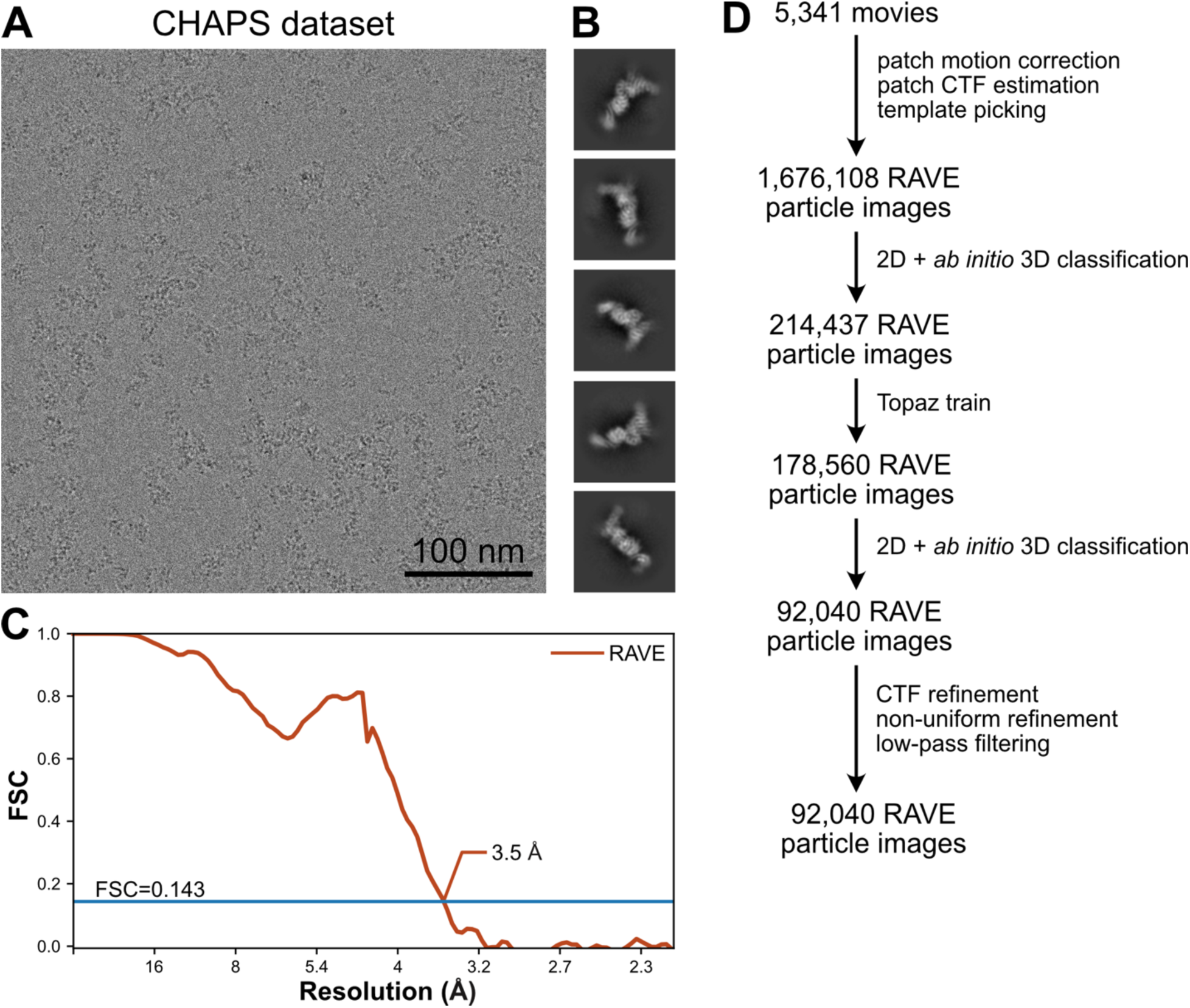
Cryo-EM of the RAVE complex. Example micrograph (**A**) and 2D class average images (**B**). **C,** Fourier shell correlation curve, corrected for masking, after a gold-standard refinement. **D,** Workflow to obtain the map of the RAVE complex.

**Figure S6.**
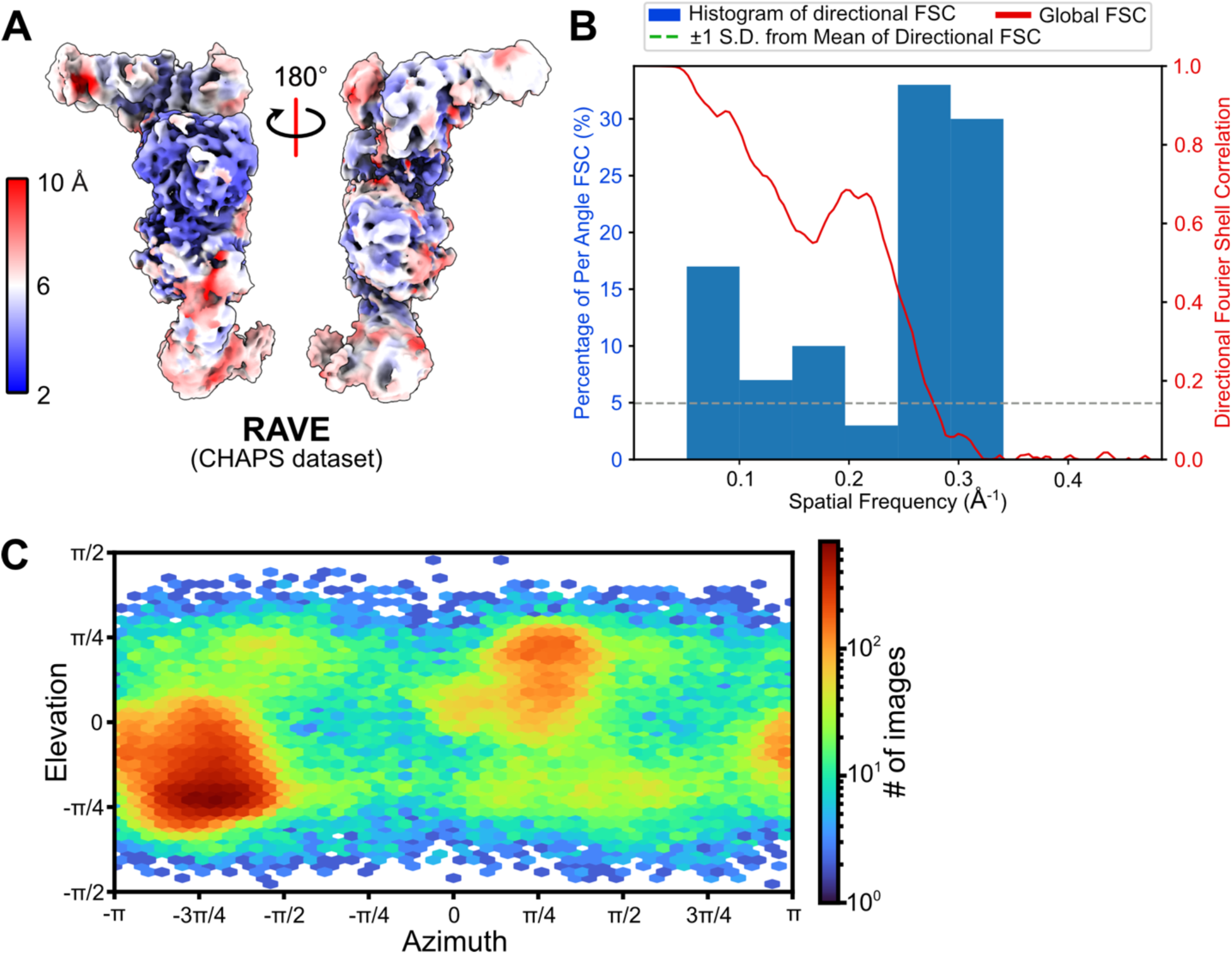
Local resolution and particle orientation distribution for the RAVE complex. **A**, Local resolution map for RAVE. **B,** Three-dimensional Fourier Shell Correlation plot for RAVE alone. **C,** Orientation distribution plot for RAVE.

**Figure S7.**
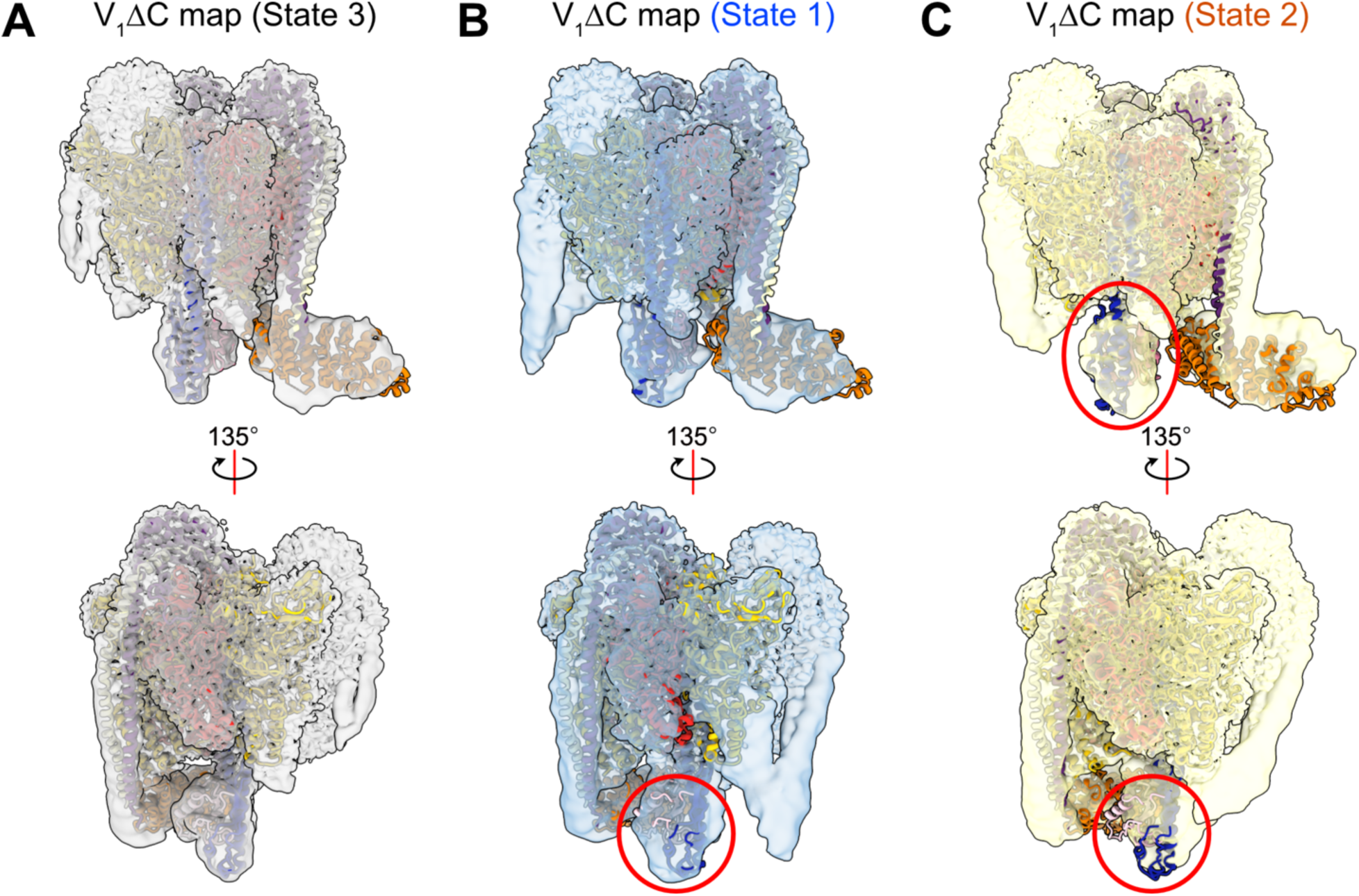
Identification of the rotational state of V_1_ΔC in the RAVE:V_1_ΔC structure. **A**, The density map of V_1_ΔC in rotational State 3 (EMD-25999) fits V_1_ΔC in the RAVE:V_1_ΔC structure with high fidelity. **B,** The density map of V_1_ΔC in rotational State 1 (EMD-25997) does not fit V_1_ΔC in the RAVE:V_1_ΔC structure. **C,** The density map of V_1_ΔC in rotational State 2 (EMD-25998) does not fit V_1_ΔC in the RAVE:V_1_ΔC structure. Regions that show poor fit are highlighted with red circles.

**Figure S8.**
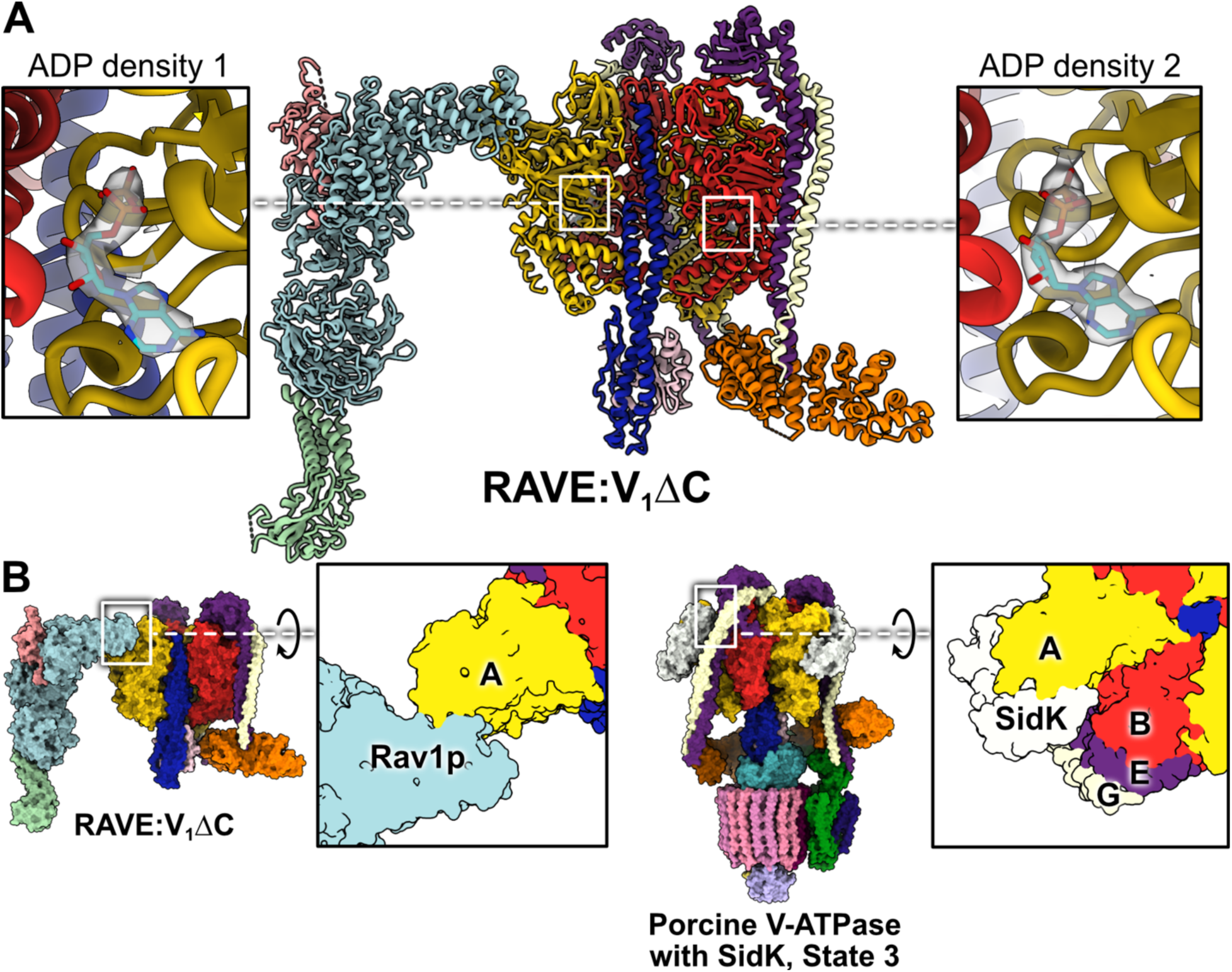
Features of the RAVE:V_1_ΔC structures. **A**, Two ADP densities are found in the RAVE:V_1_ΔC map. **B,** Comparison of RAVE binding site on yeast subunit A and SidK binding site on porcine subunit A (PDB 7U8R).

**Table S1.** Cryo-EM and atomic model building statistics.

|  | RAVE:V<sub>1</sub>ΔC<br>(EMD-45788; PDB<br>9COP) | Locally refined RAVE<br>from RAVE:V<sub>1</sub>ΔC<br>(EMD-45788) | RAVE (anisotropic)<br>(EMD-45789) |
| --- | --- | --- | --- |
| Data collection & Processing |  |  |  |
| Magnification | 75,000 | 75,000 | 75,000 |
| Voltage (kV) | 300 | 300 | 300 |
| Electron exposure (e<sup>-</sup>/Å<sup>2</sup>) | 42 | 42 | 40 |
| Exposure rate (e<sup>-</sup>/pixel/s) | 5.2 | 5.2 | 5.6 |
| Exposure time (s) | 7.7 | 7.7 | 7.5 |
| Defocus range (μm) | 0.5-2.3 | 0.5-2.3 | 0.5-2.3 |
| Pixel size (Å) | 1.03 | 1.03 | 1.03 |
| Symmetry imposed | C1 | C1 | C1 |
| Final particle images (no.) | 71,341 | 71,341 | 92,040 |
| Map resolution (Å) | 2.7 | 3.2 | 12 |
| FSC threshold | 0.143 | 0.143 | 0.143 |
| Map resolution range (Å) | 2.4 – 7.8 | 2.5 – 8.7 | 3.5 - 12 |
| Refinement |  |  |  |
| Initial model used (PDB code) | 7TMQ |  |  |
| Model resolution (Å) | 2.9 |  |  |
| FSC threshold | 0.5 |  |  |
| Map sharpening *B* factor (Å<sup>2</sup>) | locally sharpened |  |  |
| Model composition |  |  |  |
| Non-hydrogen atoms | 36501 |  |  |
| Protein residues | 5041 |  |  |
| Ligands | 4 |  |  |
| R.m.s deviations |  |  |  |
| Bond lengths (Å) | 0.007 |  |  |
| Bond angles (°) | 0.627 |  |  |
| Validation |  |  |  |
| MolProbity score | 1.68 |  |  |
| Clashscore | 6.56 |  |  |
| Poor rotamers (%) | 0.00 |  |  |
| Ramachandran Plot |  |  |  |
| Favored (%) | 95.38 |  |  |
| Allowed (%) | 4.60 |  |  |
| Disallowed (%) | 0.02 |  |  |

